# Midfrontal theta phase coordinates behaviorally relevant brain computations during cognitive control

**DOI:** 10.1101/502716

**Authors:** Joan Duprez, Rasa Gulbinaite, Michael X Cohen

## Abstract

Neural oscillations are thought to provide a cyclic time frame for orchestrating brain computations. Following this assumption, midfrontal theta oscillations have recently been proposed to temporally organize brain computations during conflict processing. Using a multivariate analysis approach, we show that brain-behavior relationships during conflict tasks are modulated according to the phase of ongoing endogenous midfrontal theta oscillations recorded by scalp EEG. We found reproducible results in two independent datasets, using two different conflict tasks: brain-behavior relationships (correlation between reaction time and theta power) were theta phase-dependent in a subject-specific manner, and these “behaviorally optimal” theta phases were also associated with fronto-parietal cross-frequency dynamics emerging as theta phase-locked beta power bursts. These effects were present regardless of the strength of conflict. Thus, these results provide empirical evidence that midfrontal theta oscillations are involved in cyclically orchestrating brain computations likely related to response execution during the tasks rather than purely related to conflict processing. More generally, this study supports the hypothesis that phase-based computation is an important mechanism giving rise to cognitive processing.

## Introduction

In recent years, cognitive neuroscientists have increasingly relied on the analysis of neural oscillations measured by M/EEG, as they appear to be a well-suited brain phenomenon to bridge behavioral observations with the neurophysiological mechanisms of brain computation (Buzsáki and Draguhn, 2004; Wang, 2010; Cohen, 2014a). An important assumption about the role of neural oscillations that fuels current research is that they provide a temporal reference for brain computations, with narrow time windows grouping neuronal activities (e.g. neuronal spiking or sensitivity to inputs; Fries, 2005) that allow effective brain computations for cognitive functions. Evidence in favor of that claim comes notably from the phenomenon of phase-amplitude cross-frequency coupling, and from direct observation of the coordination of neuronal firing according to the timing of oscillations (Canolty and Knight, 2010; Mizuseki et al., 2009). In other words, if neural oscillations are important for neural computations that subserves cognition, the phase of those oscillations should shape the timing of computations (Maris et al., 2016; Bonnefond et al., 2017; Jensen et al., 2014).

The study of cognitive control, and more specifically, of response conflict processing, has benefited from the analytic and theoretical framework provided by neural oscillations. Conflict processing refers to the ability to select a relevant response among competing alternatives that can be automatically activated. It encompasses action selection and inhibition mechanisms and is usually studied using response conflict tasks such as the Simon task (van den Wildenberg et al., 2010). When facing response conflict, an increase in theta (∼6 Hz) narrowband power is invariably observed around midfrontal electrodes (McDermott et al., 2017; Jiang et al., 2018; Pastötter et al., 2013; Cohen and Cavanagh, 2011; Nigbur et al., 2011). Importantly, this increase in power is demonstrated to reflect non-phase locked activity, suggesting that response conflict modulates the activity of ongoing endogenous theta oscillations in the prefrontal cortex (Cohen and Donner, 2013). Midfrontal theta (also sometimes called frontal midline theta) is a robust marker of conflict detection and resolution, as conflict-related theta effects have high statistical power, show strong correlations with task performance (reaction time), and relate to diseases such as Parkinson’s and OCD (Cohen and Cavanagh, 2011; Cohen and van Gaal 2014; Cavanagh et al., 2011; Min et al., 2011). Midfrontal theta phase is also demonstrated to be relevant for conflict processing, as documented by the numerous reports of conflict-modulated changes in phase synchronization between midfrontal and prefrontal and parietal regions, which suggest the existence of a fronto-parietal network of conflict processing (Vissers et al., 2018; van Driel et al., 2015; Anguera et al., 2013; Cohen and van Gaal 2013; Cohen and Ridderinkhof, 2013; Hanslmayr et al., 2008).

On the other hand, detailed predictions about the potential computational roles of midfrontal theta have remained elusive to the macroscopic scale of non-invasive human EEG, which has limited the ability to link narrowband phases to specific aspects of behavior or neural computations. One key prediction for midfrontal theta phase in conflict processing is that it provides a rhythmic structure for alternately monitoring behavior and signaling the need for control (Cohen, 2014a). In others words, this prediction implies that monitoring and signaling processes during cognitive control are more efficient at certain theta phases.

Here we provide novel evidence for a phase-specific relationship between task performance (reaction time) and midfrontal theta, using guided multivariate spatial filtering techniques to maximize the signal-to-noise ratio of single-trial data. We found that theta power-reaction time correlations are maximal at specific theta phase regions that were characteristic to each individual, and that predicted the timing of beta-band (∼20 Hz) bursts in a fronto-parietal network, as revealed by multivariate phase-amplitude coupling analyses. Contrary to our hypothesis, these theta phase-dependent effects were not specific to conditions that maximized response conflict, but rather reflected the temporal organization of a more general mechanism likely related to response execution occurring during the conflict tasks we used. All of the key findings were replicated in an independent dataset (Gulbinaite et al., 2014). The two datasets were acquired using different conflict tasks (Eriksen flanker task and Simon task; total N = 62) in two different research centers, supporting the interpretation that these findings reflect general neural signatures of cognitive control over behavior.

## Methods

We separately applied the same analysis methods to two existing datasets from studies using response conflict tasks. Here we briefly describe the key features of these two tasks; further details can be found in Cohen (2015) and Gulbinaite et al. (2014).

### Participants

30 students from the University of Amsterdam volunteered in *study 1* (Cohen, 2015). Two datasets were excluded because of excessive EEG artifacts. 34 students from the University of Groningen participated in *study 2* (Gulbinaite et al., 2014). In both studies, participants had normal or corrected-to-normal vision. The experiments were conducted in accordance with the declaration of Helsinki and with approval of local ethics committees. Written informed consent was obtained from all participants prior to the experiments.

### Task design

*Study 1* used a modified version of the Eriksen flanker task (Appelbaum et al., 2011) in which participants had to respond to a central letter while ignoring surrounding letters that acted as distractors. The strength of response conflict was manipulated in three different types of trials (Figure 1A): (i) without conflict (the target and surrounding letters were identical), (ii) partial conflict (only one side of the target had a different set of letters), and full conflict (all surrounding letters differed from the central letter). Each participant underwent approximately 1500 trials. Stimulus display lasted for 200 ms and participants had 1200 ms to respond using the left or right thumb. The two stimulus letters were mapped on two responses buttons. The next trial occurred after a 1200 ms inter-trial interval.

**Figure 1:**
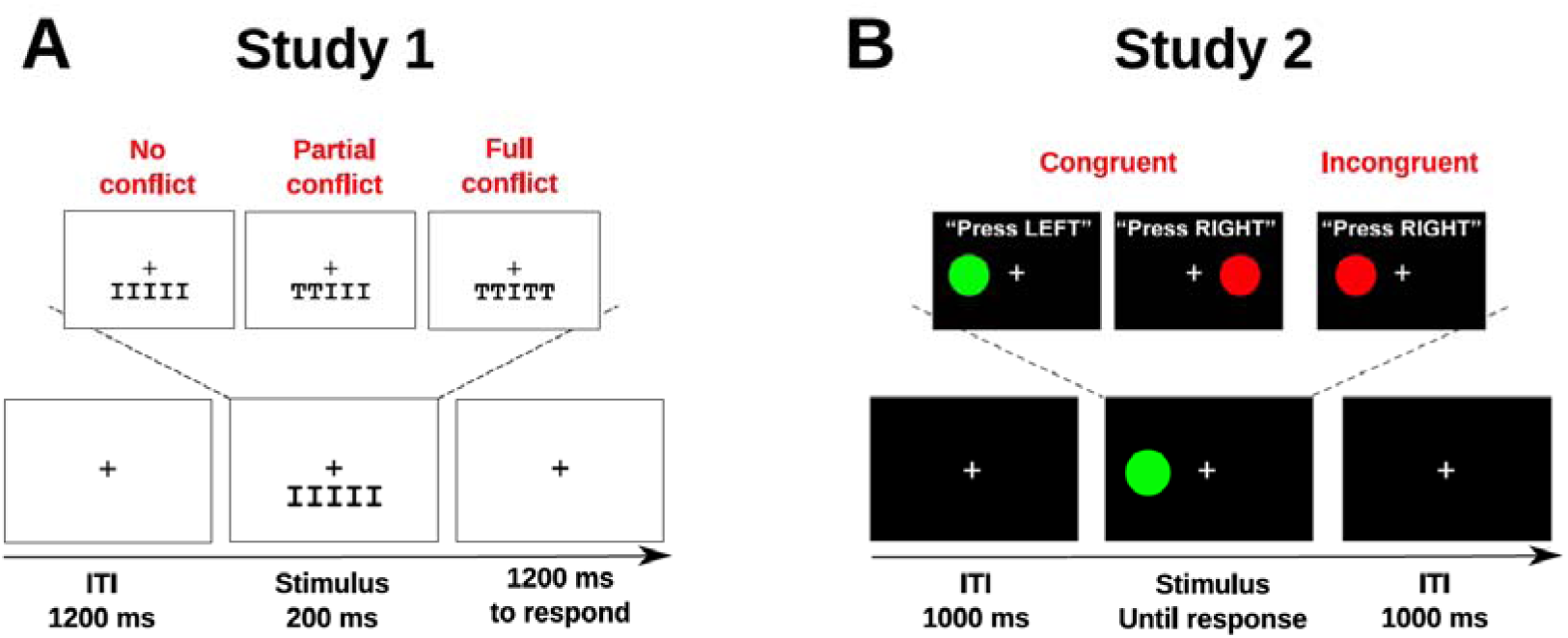
Experiment designs. A: Eriksen flanker task used in study 1. Stimuli could induce no conflict (all the letters were identical), partial conflict (letters on the left or right side differed from the target central letter), or full conflict (all surrounding letters were different from the target letter). B: Simon task used in study 2. Trials could either be congruent (color and location of the stimulus indicate the same response) or incongruent (color and location indicate different responses, hence, response conflict occurred).

*Study 2* used a Simon task with color circles as stimuli. Participants had to press a left or right button according to the color of the stimulus while ignoring its position. On congruent trials (i.e. without conflict), color and location of the stimulus indicated the same response. On incongruent trials (i.e. conflict trials), color and location indicated opposite responses. Stimuli were displayed until response or after a 1500 ms delay. After response, a fixation cross was displayed for 1000 ms before the start of the next trial. The task comprised 70 practice trials and 1024 experimental trials with the same number of possible congruency sequences: cC (congruent following congruent), cI (incongruent following congruent), iC (congruent following incongruent), iI (incongruent following incongruent).

### EEG acquisition and preprocessing

In *study 1*, EEG data were recorded at 512 Hz from 64 electrodes placed according to the international 10-20 system. Data were referenced offline to linked-earlobes, high-pass filtered at 0.5 Hz, and then epoched from −1 s to 1.5 s locked on stimulus onset. Additional electrodes were used on both earlobes and also to acquire EMG data from both thumbs’ *flexor pollicis brevis*. EMG data were used to identify small activation of the thumb muscle prior to the response. When trials contained such activation on the side of the wrong response prior to a correct one, they were labeled “mixed correct” (sometimes referred to as partial errors; van den Wildenberg et al., 2010). The identification of mixed correct trials was based on changes over time in EMG variance. The complete description of the identification algorithm can be found in Cohen and van Gaal (2014). Given that mixed correct trials are assumed to have the maximal amount of response conflict, their identification was done to label the trials in which control was most strongly implemented.

All trials were visually inspected and those containing EMG or other clear artifacts were excluded.

In *study 2*, EEG data were acquired from 62 channels at 500 Hz. The electrodes were placed according to a modified version of the 10-10 system (electrodes FT7, PO7, O1, FT8, PO8, and O2 were placed 10% below standard and electrodes F1, F2, CP1, CP2, FT7, and FT8 were not measured). Data were referenced off-line to the mastoids, high-pass filtered at 0.5 Hz, and then epoched from −1.5 s to 2 s locked on stimulus onset. Two artifact rejection steps were carried out. The first consisted in the manual rejection of trials containing clear muscle or blink artifacts and the second was an independent component analysis resulting in the subtraction of components containing eye-blinks or noise. After artifact rejection, a surface Laplacian spatial filter was applied to the data to attenuate spatial smearing as a result of volume conduction (this was done in Gulbinaite et al., 2014, to facilitate connectivity analyses).

For both studies, the epoch length was ∼2 theta cycles longer than the selected window of interest (from 300 to 1200 ms) and visual inspection of time-frequency plots did not suggest any contamination of the selected time window by the edge artifacts.

In both datasets, the ERP was subtracted from the data (separately per channel and per condition) prior to any other analysis to avoid stimulus-related phase resetting. This step removes the “phase-locked” component of the signal and ensures that the results do not reflect a stimulus-evoked transient (Cohen and Donner, 2013). This is an important methodological step that allows us to interpret our findings in terms of the phase of ongoing theta oscillations, as opposed to additive stimulus-related reset of theta oscillations. One should, however, note that there is no fully accepted method to separate the phase-locked from the non-phase-locked activity. Nonetheless, ERP subtraction is a common way of removing the phase-locked component, and has been investigated previously with regards to midfrontal theta activity (Cohen and Donner, 2013).

### Spatial filtering of the data

All signal analyses were performed in Matlab (The Mathworks, USA, version 2016a) using custom written code. Because of our a priori hypotheses regarding theta-band activity, we isolated midfrontal theta using optimal spatial filters that maximized power in the theta band. It is important to note that this approach to spatial filtering is based on statistical criteria (channel covariance matrices) and is not designed for inference about potential cortical generators. We limit our interpretations to topographical distributions to facilitate embedding in the literature. Construction of the spatial filter involves finding a set of channel weights that maximally distinguishes theta-band from broadband multichannel activity as measured through covariance matrices (Nikulin et al., 2011; de Cheveigne and Arzounian, 2015). This can be expressed via the Rayleigh quotient, in which the goal is to find a set of channel weights in vector **w** that maximizes **λ.**

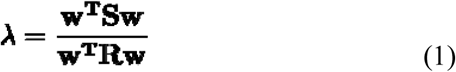

**S** is the covariance matrix derived from the data narrowband filtered in theta band (around subject-specific theta peak frequency, see below) and **R** is the covariance matrix derived from the broadband signal. The solution to equation 1 comes from a generalized eigenvalue decomposition (GED) on the two covariance matrices, defined by the equation **SW**=**RWΛ**. This method has been recommended to maximize power at low frequencies (Cohen, 2017a; Nikulin et al., 2011) and has several advantages: it increases signal-to-noise ratio, incorporates inter-individual topographical differences, and avoids the bias related to electrode selection. The column of **W** with the highest associated generalized eigenvalue is the spatial filter that maximizes the difference between **S** and **R**. This spatial filter was then applied to the data as **y**=**w**^T^**X** where **X** is the channels-by-time data matrix, and **w** is the column of **W** chosen using a method specified in the next paragraphs. Analyses are then applied to the data (component time series) in the row vector **y**.

The peak frequency of the temporal filter (for the **S** covariance matrix) was defined subject-wise by searching for the frequency of peak power in the theta band in conflict conditions at electrode FCz (conflict-related increase in theta activity has been consistently described at this location; Cohen, 2014a). To this end, we performed a time-frequency decomposition of the data by complex Morlet wavelet convolution, implemented by multiplying the Fourier transform of the EEG data by the Fourier transform of a set of complex Morlet wavelets *e^i2^*^π^*^ft^e^−t²/(2s²)^*, and taking the inverse fast-Fourier transform of the result. In the Morlet wavelets *e^i2^*^π^*^ft^e^−t²/(2s²)^*, t is time, f is frequency (ranging from 1 to 50 Hz in 60 logarithmically spaced steps), and s is the width of each frequency band, which is defined by *n*/(2π*f*) (with n logarithmically increasing from 4 to 12). Frequency specific power at each time point (t) was defined as the the squared magnitude of the resulting analytic signal (Z) as real[Z(t)²] + imaginary[Z(t)²]. Power was baseline-corrected using a decibel (dB) transform: dB power = 10*log10(power/baseline), with baseline power defined as the average power across all conditions from −500 to −200 before stimulus onset. The peak frequency was then defined within a time-frequency window of 0 to 800 ms and 3 to 10 Hz. We chose a range of 3 to 10 Hz to account for the subjects’ variability in theta peak. This peak frequency was then used as a center frequency for a Gaussian narrow-band filter (FWHM = 3 Hz) to the broadband data.

The resulting theta-filtered channel data was used to compute the **S** covariance matrix, defined in a 1000 ms (thus including several theta cycles) time window around the time of theta peak power. The same time window was used to compute the **R** covariance matrix using broadband channel data. The two covariance matrices then were used in GED to construct spatial filters. Selection of the best spatial filter in the resulting eigenvector matrix **W** was a two-step process. First, the 15 spatial filters with the highest eigenvalues were applied to the temporally unfiltered data resulting in a set of 15 components. Only those without eye-blink activity were retained. Components with eye-blink activity were identified as those with higher averaged activity at electrodes near the eyes (FP1, FPz, FP2) than around midfrontal electrodes (FC1, C1, CP1, FCz, Cz, CPz, FC2, C2, CP2, for study 1; FC1, C1, FCz, Cz, FC2, C2, for study 2). Then, the topography of the spatial filters’ activation pattern were inspected by plotting the topographical map (using the topoplot function of the EEGlab toolbox; Delorme and Makeig, 2004) of the activation pattern of the filter (**Sw**; Haufe et al., 2014). Activation patterns should be centered on the FCz electrode according to the conflict-related modulation of theta power reported in the literature (Cohen, 2014a). Thus, the component with an activation pattern consistent with midfrontal theta activity was chosen and was then analyzed using time-frequency decomposition methods (Cohen, 2017a). In 74.2% of cases, the selected component was the one associated with the largest eigenvalue. In other cases, the second or third largest component that had a midfrontal topography (monopolar with a spatial maximum around FCz) was selected instead. When several spatial filters with midfrontal topography were encountered (4 subjects), the one with the largest eigenvalue was always selected.

In this study, Morlet wavelet convolution was only used as a first step to identify the theta peak frequency for each subject and to inspect the overall time-frequency power maps of the theta component.

### Within subject brain-behavior relationships

Brain-behavior relationships across trials were investigated by correlating power in the theta band with reaction time (RT) for each participant. After filtering the previously selected component around the subject-specific peak theta power frequency found at electrode FCz during conflict, the Hilbert transform was applied to compute the analytic signal from which power was derived (according to the same squared magnitude method as reported above). A condition-specific Spearman correlation coefficient was calculated between theta power at each time point and RT resulting in a time series of correlation coefficient for each condition and each participant. Significance was evaluated using a nonparametric permutation test of the null hypothesis that the probability distribution of the participant-specific power-RT correlations is symmetric around 0. At each of the 1000 iterations, the time course of correlation coefficients was multiplied by −1 for a random subset of participants and then averaged over participants in order to generate a matrix of 1000 subject-averaged time courses under the null hypothesis. Then we subtracted the iteration averaged time course from the real subject averaged time course of correlation coefficients and divided the result by the standard deviation of the iteration averaged time course. The resulting z-valued time course was then thresholded at p<0.05. Multiple comparisons were addressed by the means of cluster-based correction so that any cluster equal or larger than 99.5% of the distribution of null hypothesis maximal cluster sizes was considered significant. This analysis served two purposes: (i) replicate correlation patterns usually observed between theta power and RT during conflict and (ii) define a time window of interest (in which correlations were significantly different from 0) for phase-resolved analyses.

### Within subject phase-resolved brain-behavior relationships

In order to investigate if the theta power-RT correlations depended on theta phase, we computed theta power-RT correlation coefficient according to the different possible phase values in 20 discrete phase bins. To this end, the phase angle time series of the previously Hilbert-transformed theta-defined component of each trial was divided in 20 linearly spaced phase bins with values ranging from −π to π. For each theta phase bin, the corresponding power was extracted and averaged for that trial. For instance, the different power values extracted when phase equals π/2 across the phase angles time series were averaged in order to get the mean power at the π/2 phase for that trial (Figure 2 B and C). This operation was repeated for all trials and separately for each condition. Phase-specific mean power was then correlated with RT, resulting in 20 power-RT coefficients for each condition and subject (Figure 2 D). These phase-specific correlation coefficients were then averaged across subjects for each condition (Figure 2 E).

**Figure 2:**
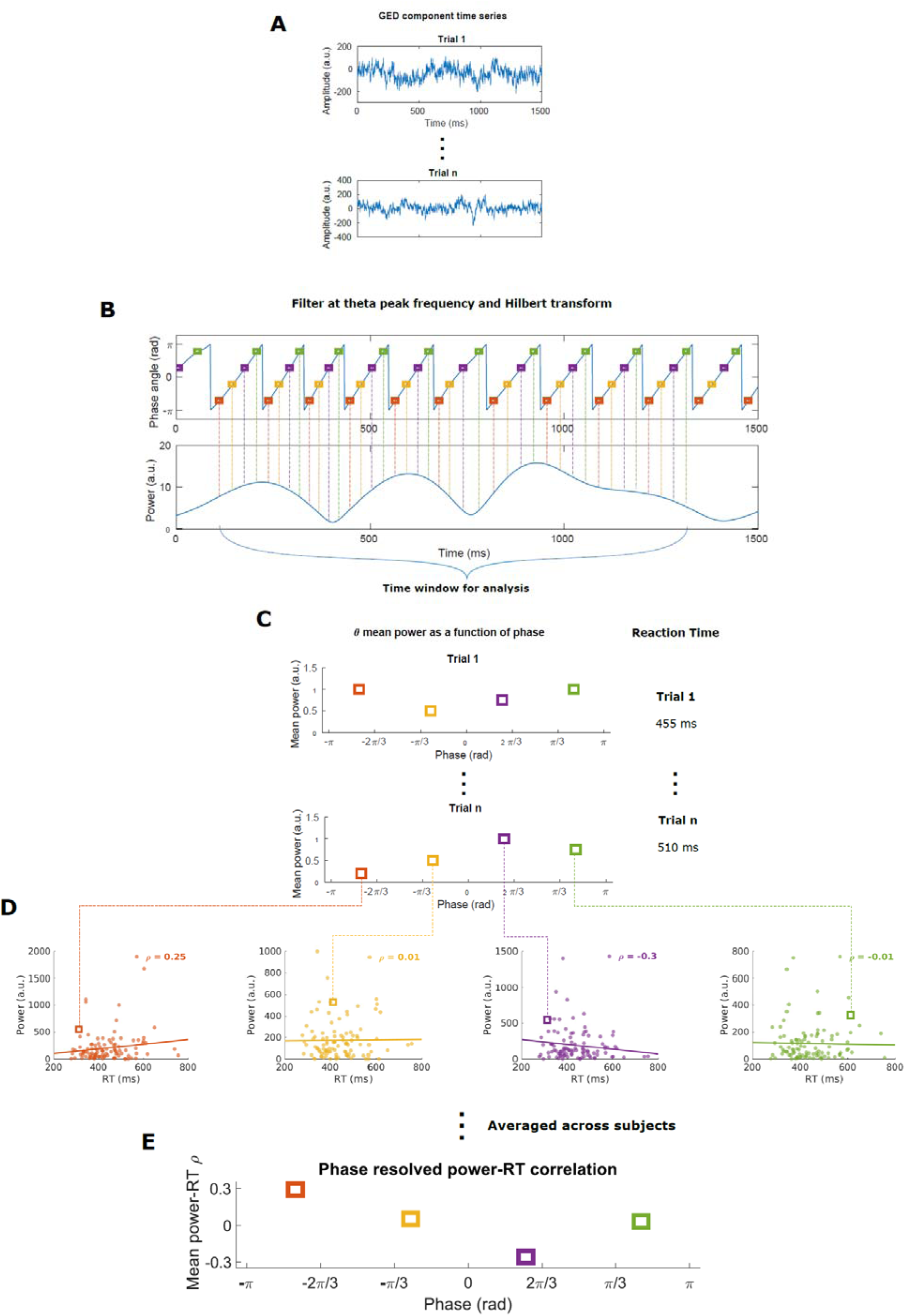
Overview of the method used to compute phase-resolved power-RT correlations. Each trial (A) was filtered using the theta frequency of maximum power previously found at FCz, and the Hilbert transform of that filtered signal was taken to extract phase and power. Power at each corresponding theta phase bin was extracted (B) and the average power for each phase bin was computed for all trials (C). Then, Spearman’s ρ between power and RT were calculated at each phase bin for all subjects (D) and were then averaged across subjects (E).

### Within subject phase-resolved power-RT correlation analyses

In order to test whether power-RT correlations significantly varied according to phase, we fitted the correlation values to sine waves for each subject and condition. These analyses were done using R (version 3.4.2; R core team, 2017) and the {minpack.lme} package for nonlinear least square fitting (Elzhov et al., 2016). Data were fitted by applying the nlsLM function to the model: ρ∼*A\**sin(2*π*\*F*\*phasebin+θ), where ρ is the correlation coefficient, and *A*, *F* and θ, are the amplitude, frequency, and phase parameters to estimate, respectively. Parameter estimates were then used in a harmonic addition of a sine and a cosine (which is equivalent to a single sine wave allowing for a phase shift) to enter in a linear model following: ρ ∼(*A*sin(2π*F*\*phasebin+ θ) + *A*cos(2π*F*\*phasebin+ θ)). This was done in order to estimate the overall significance and the goodness of fit of the estimated sine parameters.

The amplitude parameter and the model R² were compared at the group-level between all conditions using wilcoxon paired tests with a Bonferroni corrected threshold of p < 0.05/10 = 0.005 because of the 10 possible conflict condition comparisons. We also statistically tested the null hypothesis that the population distribution of preferred phases is uniform on the circle using Rayleigh’s Z.

### Cross-frequency coupling

We next sought to investigate whether higher-frequency dynamics were locked to midfrontal theta phase in a manner relevant to the phase of peak power-RT correlations. Assessing the modulation of higher-frequency power by lower-frequency phases is usually performed by means of a phase-amplitude coupling index. However, here we chose a different approach that would take into account the results of theta phase-resolved power-RT correlations. We used GED (Cohen, 2017a) to define a component that maximizes activity around the phase of maximum brain-behavior correlation. For each subject, the **R** matrix was defined as the data filtered between each subject theta frequency + 2 Hz and 80 Hz. This frequency range was selected to eliminate both theta activity and higher-frequency activity unlikely to reflect brain dynamics. For the **S** matrix, the same filtering was applied but we extracted the data around the empirically determined phase of maximum theta power-RT correlation. We identified each occurrence of the preferred theta phase in each trial (between 300 and 1200 ms), then extracted the data from ⅛ of a theta cycle before that phase to ⅛ of a cycle after that phase (the exact number of ms depended on subject-specific theta frequency); the covariance matrices from these time windows were averaged to create the **S** covariance matrix. The spatial filter with the largest generalized eigenvalue was selected to define the component. In this case, rather than isolating narrowband theta dynamics, GED served the purpose of enhancing the signal’s characteristics in higher frequencies occurring around the phase of maximum theta power-RT correlation.

Next, we applied time-frequency decomposition on the selected component to compute power for each frequency (from 10 to 50 Hz) resolved according to the phase of the GED-identified theta component which was divided into 51 bins (we used more bins than for phase-resolved power-RT correlation to have a better phase resolution for the phase-frequency power maps). We used the same Morlet wavelet convolution methods as described above. Each phase-frequency map was phase-shifted so that phase = 0 corresponds to each subject’s phase of maximum theta power-RT correlation. Then, decibel-transformed power (computed using the same methods and baseline as described above) was z-scored for each frequency in order to quantify phase-specific modulation of power. Finally, clusters of significant phase-specific changes in power were defined using permutation testing. For each iteration, all the data after a randomly-defined relative theta phase (between the 10th and the 40th bins) were shifted and a difference map between the phase-shifted map and the original unshifted map was calculated. This operation was carried out 10,000 times, generating a distribution of difference maps under the null hypothesis. The average real data map was z-scored using the mean and the standard deviation of the difference maps. The resulting map was then thresholded at p<0.001. Multiple comparisons were addressed by the means of cluster-based correction so that any cluster equal or larger than 99.9% of the distribution of null hypothesis cluster sizes was considered significant. Power data from the biggest significant cluster was then extracted and compared at the group level between conflict conditions using a one-way ANOVA.

## Results

### Behavioral results

The classical slowing of RTs as a function of conflict was observed in both studies (study 1: F(1, 26) = 6.17, p = 0.016; study 2: F(1,32) = 20.81, p < 0.001). Conflict trials were also significantly less accurate in study 1 (study 1: F(1, 26) = 60.6, p<0.0001), and showed a similar trend in study 2 (F(1,32) = 3.54, p = 0.069). For a more detailed description of behavioral results, please refer to Cohen (2015) and Gulbinaite et al. (2014).

### GED results, component topography, and time-frequency power

Given our a priori focus on the theta band, we applied GED to find a spatial filter that best isolates theta band activity from the broadband activity in the signal. Data reduction using GED was implemented using the theta-filtered channel covariance matrix data as the **S** matrix (mean theta frequency used for filtering: study 1: 4.6 ± 1.6 Hz, study 2: 5.4 ± 0.7 Hz) and the broadband data channel covariance matrix as the **R** matrix. Figure 3 A and D show the topography of the activation pattern of the selected spatial filters averaged across participants. For both studies, a clear higher activation can be seen around midfrontal electrodes, consistent with the topography of what is usually reported in the literature as midfrontal theta. Time-frequency decomposition of the selected component in study 1 shows a clear increase in power compared to baseline in the theta band (Figure 3B). This power increase was stronger with increasing conflict strength and was the highest for errors (Figure 3C1). Figure 3C1 and C2 shows the same pattern focusing on the power time course of the peak theta frequency and the peak of power for that frequency respectively. In study 2, the same results were observed with a clear theta power increase that was stronger in the situations of strongest conflict (cI trials) compared to all other trials (cC, iI, and iC trials; Figure 3E, F1 and F2). As in study 1, errors showed the strongest increase in theta power. As a whole, these results show that applying GED for spatial filtering successfully produced a signal that exhibits all effects that are typically reported in conflict resolution studies. Given that the data in study 2 were Laplacian-transformed (which was not the case in study 1), we Laplacian-transformed the data from study 1 and recomputed the GED spatial filters as well as the time-frequency power maps to ensure that the main results did not depend on Laplacian transformation. Applying the Laplacian transform did not change the results both at the subject-or at the group-level with similar topographies and nearly identical time-frequency power maps (see supplemental figure S1).

**Figure 3:**
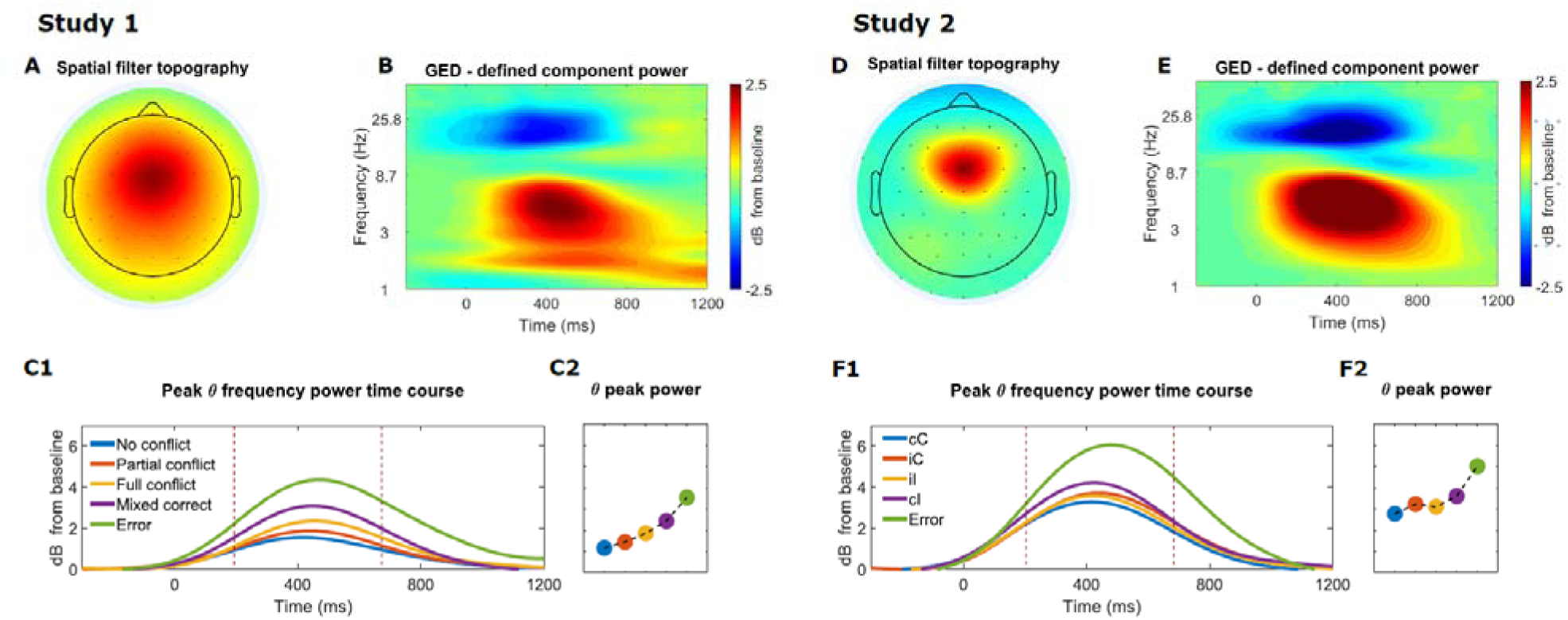
A: Topographical map showing the activation pattern of the spatial filter designed by GED in study 1. B: Time-frequency plot of power (decibel-transformed) of the component defined by GED in study 1. C1 and C2: theta power time courses (vertical dashed lines indicate the time window selected to compute the peak of power) and peak of power according to the strength of conflict. D to F2: same plots for study 2.

### Theta power-RT correlations

Following previous reports (Cohen and van Gaal, 2014), we first investigated brain-behavior relationships by means of cross-trial correlations between power and RT (ignoring theta phase). This analysis also served the purpose to define a time-window of significant theta power-RT correlation to investigate the potential modulation according to theta phase. For each subject and each condition, Spearman’s rho was computed across trials between power at the peak theta frequency and RT at each time point, resulting in one time course of power-RT rho values per subject and condition (see Figure 4, for group average result). In both studies, an increase in theta power-RT correlation began around 200-300 ms, peaked around 500-600 ms and was immediately followed by a decrease in all conditions, except error trials. The pattern of power-RT correlation for errors was clearly distinct, beginning with a sharp decrease around 200-300 ms, followed by an increase peaking around 600-700 ms and finally a return to zero. Using cluster-based permutation tests, we tested the null hypothesis that the population distribution of the power-RT correlations is symmetric around zero. Rejection of the null hypothesis at p < 0.05 was observed from roughly 300 ms to 1200 ms in Study 1 (Figure 4A), and from 200 ms to 1200 ms in Study 2 (Figure 4C). Based on this result, for subsequent phase-resolved analyses we selected the following time windows: 300 to 1200 ms for study 1 and 200 to 1200 ms for study 2.

**Figure 4:**
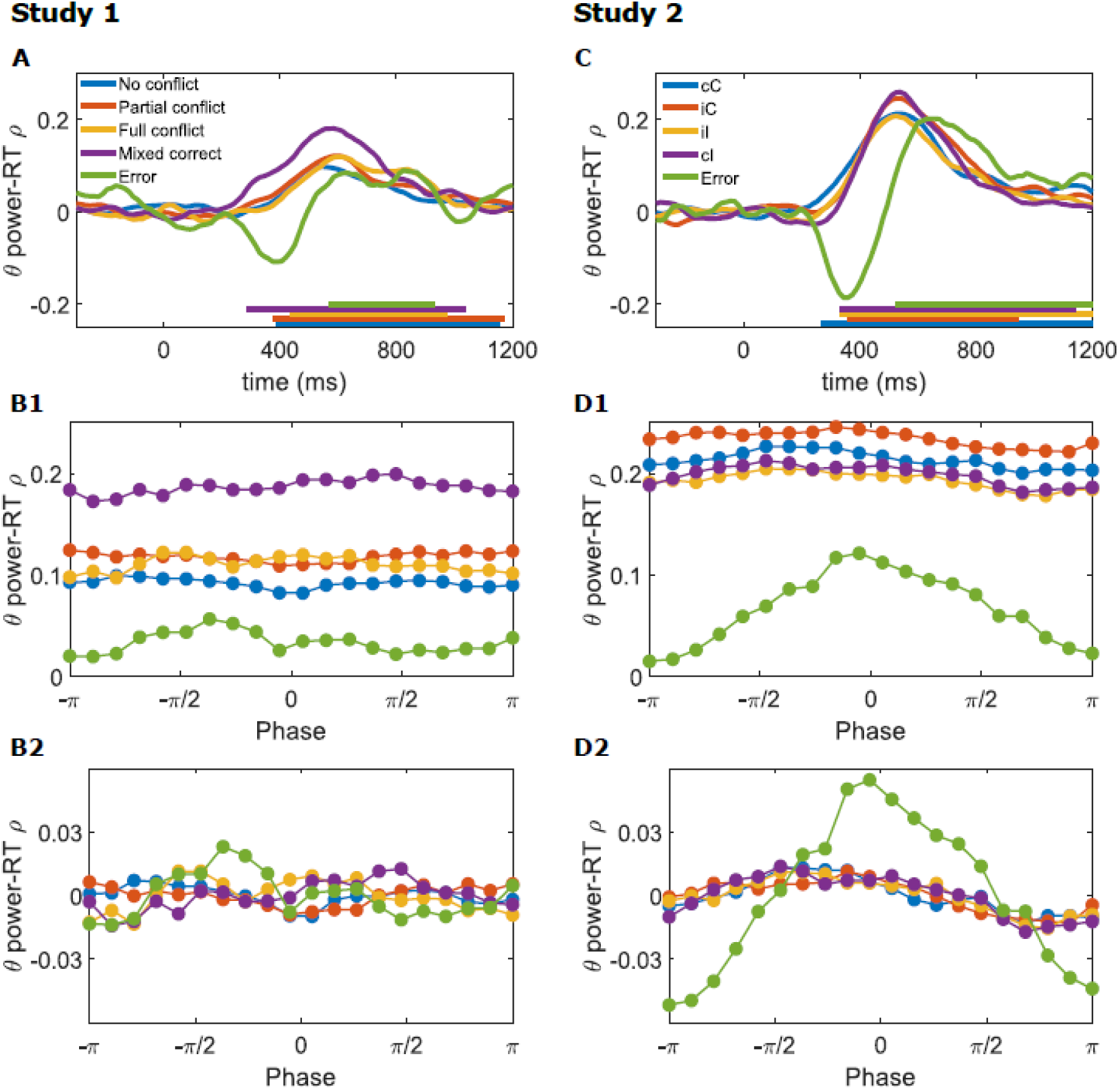
Time course of Spearman’s ρ between theta power and RT averaged across subjects for each condition in study 1 (A) and 2 (C). Horizontal bars above the x-axis denote windows of statistical significance at the group-level. B1 and B2: phase-resolved Spearman’s rho between theta power and RT. These plots show the average (B1) and detrended (B2) theta power-RT correlation coefficient according to theta phase. D1 and D2: same plots for study 2.

In both studies, and consistent with previous reports (Cohen and Cavanagh, 2011; Cohen and van Gaal, 2014), the highest peak of correlation was observed for the trials with the strongest conflict (mixed correct and cI trials respectively). However, statistical testing confirmed this was the case only in study 1. We extracted the peak of maximum correlation from the time windows defined above and applied pairwise Wilcoxon paired test to compare between conflict conditions. We found that partial errors were indeed the situation in which RT-power correlations were maximal (mixed correct vs. no conflict: p = 1.26*10^−6^; partial error vs. partial conflict: p = 7.2*10^−5^; full conflict vs. no conflict: p = 0.0014; all other conflict conditions comparisons were non-significant at p>0.005 using Bonferroni correction). However, in study 2, the peak of maximum correlation did not differ between conflict conditions (all p>0.005).

### Phase-resolved theta power RT correlation

We further investigated the influence of theta phase on brain-behavior relationships. We reasoned that if theta phase defines certain time-windows of behaviorally-relevant brain computations, fluctuations of theta power-RT correlations would vary as a function of theta phase. To test this, we averaged power in 20 discrete theta phase bins (in previously defined significant theta power-RT correlation time windows), and correlated this phase-resolved average theta power with RTs. This was done separately for each subject and condition. Given that the size of the selected time windows was 900 ms for study 1 and 1000 ms for study 2, and given that the average theta frequency was 4.6 ± 1.6 Hz in study 1 and 5.4 ± 0.7 Hz in study 2, all phase-resolved theta power-RT correlations were obtained based on at least 4 theta cycles.

At the group-level, correlations were higher for some theta phases than for the others in all conditions and in both studies. This can be seen in Figure 4B1 and D1, but is more evident in the detrended coefficients in Figure 4B2 and D2. Closer inspection, however, revealed subject-level phase preferences across conditions and subjects (see sine fit plots in Figure 5A). We therefore computed nonlinear least square estimations for each subject and for each condition, specifying a sine function with sine parameters to be estimated from the data. The estimated parameters were then entered in a linear model to inspect overall significance and the goodness of fit of the models.

**Figure 5:**
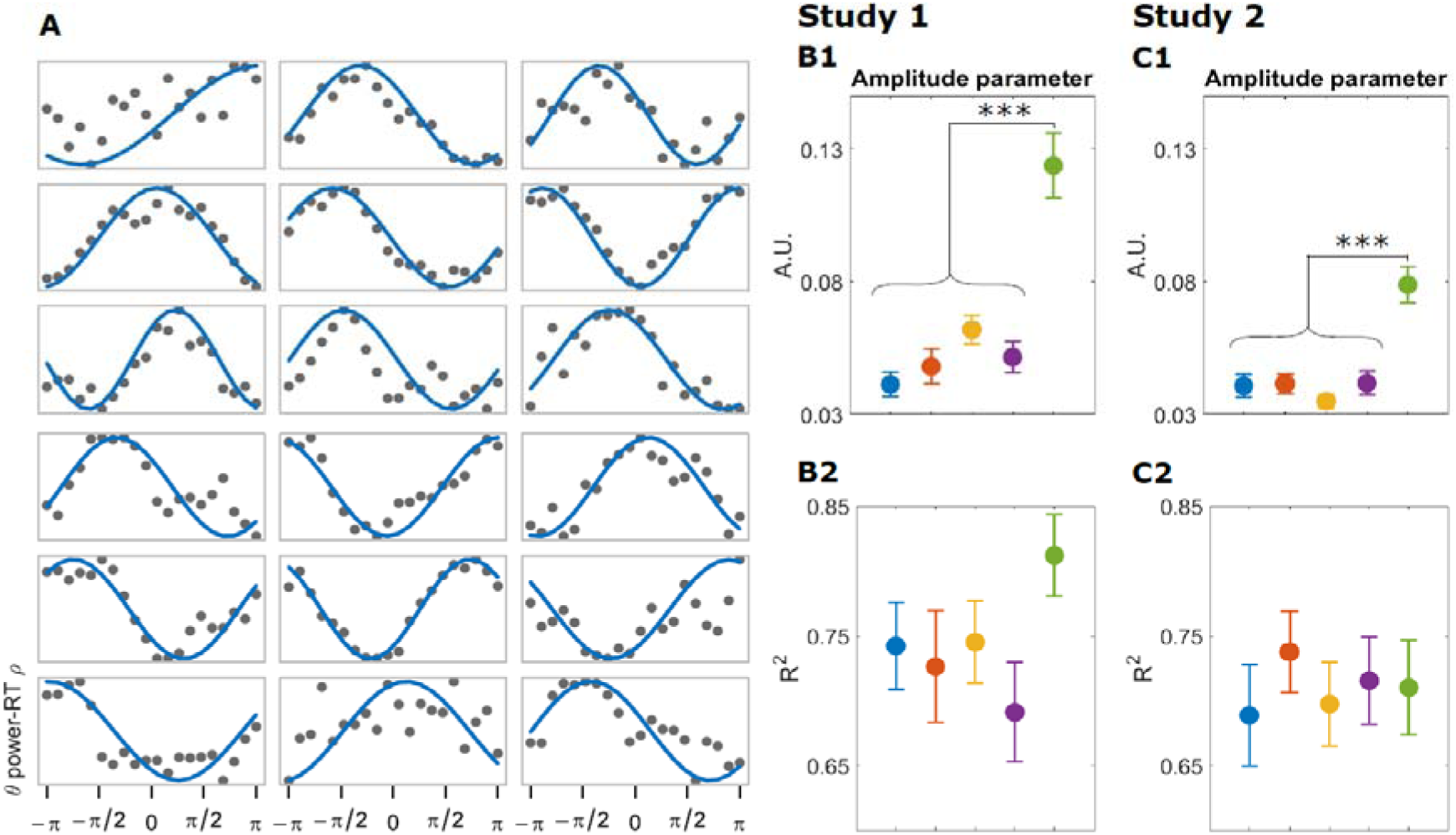
A: example of the sine fit for 18 subjects in the no-conflict condition in study 1. All sine fit plots can be seen in supplementary figure S2 and S3. B1 and C1: average amplitude parameter according to conditions for study 1 and 2, respectively. In both studies, errors had a significantly higher amplitude parameter than all other conditions (p<0.001). B2 and C2: average R-squared according to condition for study 1 and 2 respectively. Color coding of conditions is the same as in Figure 4 (weaker to stronger conflict and errors from left to right). Error bars represent the standard error of the mean.

#### Sine fit

All sine fit graphs are depicted in supplementary figure S2 and S3, and an example fit can be seen in figure 5A. For each condition and in both studies, sine waves were significant predictors for the observed phase-resolved power-RT correlations, across subjects and conditions in both studies. Since the amplitude parameter is the best predictor of the size of the phase-resolved effect, we only report these results here (see supplementary table S1 to see the proportion of subjects with significant parameters r). In study 1, 100, 96, 96, 100 and 100% of the nls fits showed significant amplitude parameters for the no conflict, partial conflict, full conflict, mixed correct, and error conditions respectively. In study 2, 100, 97, 100, 97, 97 % of the nls fits showed significant amplitude parameters for the cC, iC, iI, cI and error conditions respectively. Overall, for both studies, 95.8% of the linear models using nls-defined parameters significantly fitted the data. This indicates that in both studies, and in all conditions, theta power-RT correlations were stronger for some phases than for others.

#### Group-level amplitude and R-squared comparison

We tested whether the theta phase influence on brain-behavior relationships depended on the strength of conflict, assessed as the amplitude of the fitted sine wave, as this indicates the magnitude of theta phase on theta power-RT correlations. We performed pairwise Wilcoxon tests on the condition-specific amplitude parameters, using a significance threshold of p < 0.005 (Bonferroni correction for 10 comparisons). As can be seen in figure 5B1 and C1, the amplitude parameter was not statistically different across correct conditions, and was different for error trials. Indeed, amplitude for errors was significantly higher than for other conditions in both studies (all error-other condition pairwise wilcoxon paired test were significant at p<0.0001 in both studies; all other pairwise comparisons were non-significant). We subsequently tested whether the strength of conflict impacted the fit of the models to the data (R²). In both studies, R² did not significantly differ between conditions (all pairwise wilcoxon paired test were non-significant) (Figure 5B2 and C2), suggesting that the goodness of fit of the models were similar across the different levels of conflict in both studies.

#### Phase consistency

Given the fluctuations of theta power-RT correlations according to theta phase, we also investigated whether the phase preference (the phase of maximal theta power-RT correlation defined by the empirical data) was consistent across subjects in each condition. To that end, we computed phase clustering of the phase of maximal theta power-RT correlation for each condition (see supplemental figure S4). Preferred phase was not clustered and was relatively homogeneously distributed, except for errors in study 2 which showed significant clustering (p =0.002) around phase = 0. In all other cases, clustering measures were non-significant, confirming that the phase of maximum theta power-RT correlation was subject- and condition-specific.

#### Power and RT as a function of phase

To further contextualize the theta-phase-modulated brain-behavior correlations, and to evaluate whether the fluctuating correlations were due to information fluctuations in the individual variables, we examined whether theta power or RT independently were modulated by theta phase. We used the same theta phase discretization as for the correlation analyses, but now focusing on theta power and RT separately rather than on their correlation. Supplementary figure S5B1-D1, and S5B2-C2 show the detrended average theta power and standard deviation, respectively, according to phase for study 1 and 2. Theta power and its standard deviation in both studies displayed a clear modulation according to phase, thus indicating that power was stronger at certain theta phases and lower at others. Such asymmetries in power over phase bins also point to a non-sinusoidal waveform shape (Cole and Voytek, 2017). Conversely, average RT and its standard deviation in both studies (supplementary figure S5F1-H1 and S5F2-H2) showed no clear evidence of phase-dependent fluctuations.

### Control analyses

Although all the results argue in favor of the hypothesis that theta phase influences brain-behavior relationships, we wanted to test whether the effect we found was specific to the theta band and whether the effect was reproducible within each dataset individually (cross-validation). We chose to test phase-resolved brain-behavior correlations in the beta band, since beta suppression is also typically observed around RT in conflict tasks (see Figure 3B and E) and since beta activity has been shown to relate to RT and errors (Doyle et al., 2005; Torrecillos et al., 2015).

#### Theta band specificity - test of beta phase-resolved power-RT correlations

We extracted power from the Hilbert transform of the beta bandpass-filtered (between 15 and 20 Hz) theta-defined component and computed power-RT Spearman correlations at each time point. We selected the beta band because it is another frequency range that is observable in midfrontal topographical regions during conflict tasks (Figure 3). As can be seen in supplementary figure S6A and S6C, the patterns of power-RT correlations are clearly different from the theta results in both studies. For all conditions in study 1, only negative correlations can be observed occurring between around 500 and 1000 ms and with an effect roughly half the size of what was observed in theta. Study 2 shows the same pattern of negative correlation occurring this time mostly between 500 and 900 ms with a significant increase beginning at 1000 ms.

In order to test theta-specificity of phase-resolved results, we computed beta power-RT correlation for the 20 beta phase bins during the significant theta power-RT correlation time window. Supplementary figure S6B1-B2 and S6D1-D2 show the average and detrended version of the phase-resolved beta power-RT correlations for study 1 and 2. Even though visual inspection did not suggest an effect of phase, given a high interindividual variability in theta phase preference when investigating theta phase-resolved power-RT correlation, we nonetheless inspected beta phase-resolved results at the subject level and performed the same sine wave modeling as for the theta results. As expected, no clear oscillation of correlation according to the beta phase was observed: sine wave model accounted for the data with 50, 39, 28, 25 and 25% of significant nls fits for the no conflict, partial conflict, full conflict, mixed correct, and error conditions respectively in study 1. For study 2, the same results were observed with 41, 29, 26, 35, and 47% of significant *nls* sine fits amplitude parameter for the cC, iC, iI, cI and error conditions respectively (see supplementary table 1 for details of all parameters). This control analysis provides further support for theta-band specificity in phase dependent power-RT correlations.

#### Split-half reliability

In order to evaluate if the phase dependent theta power-RT correlation effect was consistent across trials within subject, we re-analyzed all the data separately for two halves of the data collapsed across conditions. This operation was repeated 100 times by randomly selecting two groups of trials. In both studies, results matched the pattern of theta power-RT correlation observed when analyzing all the data. We next examined whether the preferred phase that maximized power-RT correlations was consistent between the two halves of the data. We therefore computed the subject-averaged difference in preferred phases between the two halves, for each iteration; this generated a distribution of 100 phase difference values, on which we computed phase clustering. If the preferred phases are consistent, the different distribution should cluster around 0 radians across subjects. We observed significant clustering of the phase differences around 0 for both studies (study 1: p<0.001, study 2: p<0.001; supplemental figure S7A and S7B). We further inspected the overall correlation between the two halves of the data for those 100 iterations and observed significant positive correlations in 100% of the cases in both studies (mean ρ = 0.48, mean p < 0.0001 for study 1 and mean ρ *=* 0.44, mean p < 0.0001 for study 2. Taken together, these results show that the phase effect we found was consistent across the experiment.

### Cross-frequency coupling

The previous sets of analyses demonstrated that theta phase is relevant for brain-behavior relationships. We next investigated whether this same theta phase is relevant for higher-frequency EEG activity. We focused on phase-amplitude CFC (Canolty & Knight, 2010) under the assumption that if theta phase is relevant for brain computations that serve behavior, then specific patterns of CFC could appear around the phases of maximal power-RT correlations.

We applied the GED framework to identify a component that maximizes EEG activity around the phase of maximum theta power-RT correlation at higher-frequencies. Figure 6A1 and A2 show the topography of the activation pattern of the spatial filter for study 1. Figure 6A2 is the laplacian-transformed spatial filter topography that we computed in order to better compare with the topography of study 2 (Figure 6D) in which the data were scalp-Laplacian-transformed. In both studies, but even more so in study 2, the topographies highlighted centro-parietal and parietal electrodes. In study 1 higher activation was observed around midfrontal electrodes.

**Figure 6:**
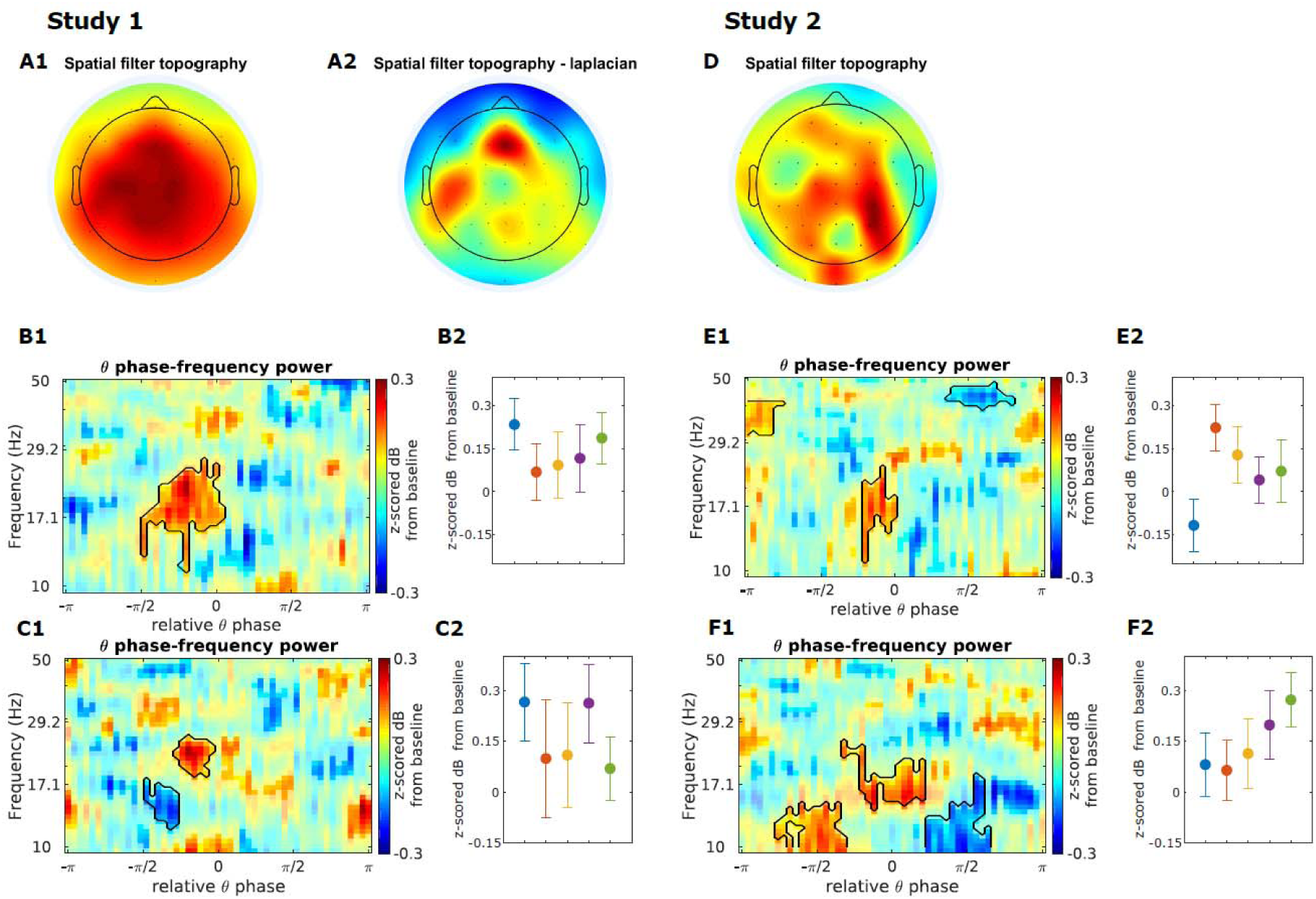
A1-A2: topography of the spatial filter defined by GED (A2 is the scalp Laplacian-transformed topography) in study 1. D: spatial filter topography for study 2. In both studies, activation around centro-parietal and parietal electrodes can be observed. More midfrontal activation is seen in study 1. B1 and E1: theta phase-frequency power map for study 1 and 2, respectively. Contour lines show significant clusters (with sizes exceeding the mean cluster size plus 2 standard deviations after cluster correction of permutation testing). B2 and E2: power extracted from the common cluster of study 1 and 2, respectively. C1 and F1: theta phase-frequency power map computed by averaging the maps calculated using a cross-validation approach from study 1 and 2, respectively. C2 and F2: power extracted from the common cluster from study 1 and 2, respectively. Color coding of conditions is the same as in previous figures and goes from weaker to stronger conflict and errors from left to right. Error bars represent the standard error of the mean.

We inspected power at each theta phase bin (51 bins) and for each frequency. Figure 6B1 and E1 show theta phase-frequency power maps for study 1 and 2 respectively. In both cases cluster-corrected permutation testing revealed significant increases in power between 0 and − π*/2* and in the upper alpha-beta frequency range (biggest cluster in the map) (power peaks were at phase −π*/5* and at 19 Hz for study 1 and at phase −π*/6* at 17 Hz for study 2). Note that these phase angles are relative to subject-specific peak power-RT correlations, not absolute theta phase angles. Power was extracted in the biggest significant cluster and was compared between conditions (Figure 6B2 and E2). In both studies, power did not differ between conflict conditions (study 1: F(4, 135) = 0.44, p = 0.8; study 2: F(4, 165) = 0.62, p = 0.65), suggesting that phase-specific increase in power reflected a general mechanism that was related to response execution but not specific to conflict processing.

To ensure that these results were robust, and that they were not biased by overfitting, we re-analyzed the data using a cross-validation approach. We used 90% of the data to compute the cross-frequency-coupling spatial filter using the same criteria as previously described, and applied it to the remaining 10% of the data. This operation was carried out 10 times, and the resulting phase-frequency maps were averaged over iterations for each subject. The same permutation testing as in the original CFC analyses was then applied. The spatial filter topographies looked the same as the ones in our first set of analyses, and thus are not depicted in figure 6. Although some differences with the first analyses can be seen, the previously described common cluster was also present here with a significant increase in power between 0 and −π*/2* in the upper alpha-beta band (Figure 6C1 and F1). In both studies the frequency and theta phase of the peak of power from that cluster was very similar to the first analysis with a peak at 22.8 Hz −π*/5* for study 1 and at 24.8 Hz and −4π*/7* for study 2. Similar to the first analysis, power extracted from those clusters did not show any significant change according to the strength of conflict (Figure 6C2 and F2; study 1: F(4, 135) = 0.48, p = 0.7; study 2: F(4, 165) = 0.87, p = 0.5). As a whole, both the cross-validation approach and the replication in the 2 independent datasets strongly suggest that there was a specific increase in power in the beta band around −π*/5* relative to the phase of maximum power-RT correlation that was not specific to the strength of conflict but rather generally involved in response execution.

## Discussion

In their formulation of the theta-gamma neural code, Lisman and Jensen (2013) emphasized that theta oscillations may “provide a way of ordering multipart messages.” Although their conclusions specifically related to the role of hippocampal theta phase in spatial location and memory, this formulation also echoes with the proposal that neural oscillations have a general role in providing time references for brain computation (Fries, 2005; Voytek and Knight, 2015). With this study we investigated a key hypothesis of one potential role of midfrontal theta oscillations, namely, whether they define specific time windows during which brain computations that are relevant for response conflict processing occur (Cohen, 2014a). We computed phase-resolved brain-behavior correlation based on a multivariate analysis method that maximizes signal-to-noise ratio of relevant features of the EEG signal. Results from two independent studies confirmed the robustness of the findings.

### Implications for the role of midfrontal theta in brain computations

Although theta phase has already been shown to be important for perceptual and motor activities (Drewes and van Rullen, 2011; Tomassini et al., 2017; Han and van Rullen, 2017), to our knowledge this study provides the first empirical evidence of a temporal organization relying on an internal reference frame defined by the phase of ongoing theta activity around midfrontal electrodes during cognitive control (Panzeri et al., 2010). It has been proposed that different computations for conflict processing would be implemented by a neural microcircuit in the midfrontal cortex in a layer-specific manner (Cohen, 2014a). Although it would be hazardous to speculate on the contribution of the layer-specific activity in the scalp EEG signal and their theta phase-relative organization, if midfrontal theta oscillations organize conflict processing, it is plausible that different theta phase regions could provide temporal constraints for computations occurring in different cortical layers. In that manner, specific computations during conflict processing may take place at specific theta phases. For instance, the relevant phase described in this study could be specific to a largely motor-related mechanism that underlies response execution, while time windows of opposite phases could preferentially support conflict integration, or communication with long range task-relevant cortical areas responsible for selective inhibition or modulation of attention. A more physiologically detailed understanding of the role of midfrontal theta oscillations in conflict processing will require additional work with invasive recordings (including local field potential and single units). However, the results of this study provide strong evidence for the oscillatory nature of the signals involved in conflict processing and, most importantly, gives insights on the existence of a temporal organization during conflict processing by midfrontal theta oscillations in humans.

Our findings also have implications for ongoing discussions about the nature of midfrontal theta. In particular, whether the EEG signature of response conflict monitoring truly reflects a “neural oscillation” or simply a non-oscillatory transient remains debated (Yeung et al., 2007, Trujillo & Allen, 2007, Cohen & Donner 2013). Our phase-dependent brain-behavior findings provide new evidence for the interpretation of a neural oscillation. A phase-specific finding that is present after removing the phase-locked part of the signal reflects amplitude modulations of ongoing narrowband activity, and thus is inconsistent with an interpretation of the conflict-modulated theta burst as an additive transient (which was removed from the data prior to our analyses).

### Midfrontal theta phase influences the strength of brain-behavior relationships

Midfrontal theta has been consistently shown to be relevant for behavior during conflict processing, as RT strongly correlates with power (McDermott et al., 2017; Jiang et al., 2015a; Jiang et al., 2015b; Jiang et al., 2018). This correlation is especially strong for high-conflict trials and immediately preceding the response (Cohen and Cavanagh, 2011; Cohen and van Gaal 2014). The present study provides yet another replication of these results (in Study 1) and, in line with our hypothesis, further shows that the strength of these correlations depends on the phase of the ongoing midfrontal theta oscillation. The effect was specific to the theta band and reliable across the experiments in two different conflict tasks (Eriksen flanker task and Simon task). Furthermore, specific investigation of phase-resolved power revealed that theta power was non-uniformly distributed around the theta cycle, which is indicative of a non-sinusoidal waveform shape (Cole and Voytek, 2017). This suggests that the influence of theta phase on brain-behavior relationships may have resulted from the waveform shape of the midfrontal theta oscillation. The phase effect on RT-power correlation was independent on the strength of conflict, which points to a general response execution mechanism occurring during conflict processing rather than a purely conflict-related mechanism as we hypothesized. We nonetheless observed some condition differences, with a greater phase effect for errors compared to all other conditions. This error-specific modulation is consistent with a general observation that errors have qualitatively distinct patterns of theta-RT correlations compared to correct responses, with negative correlations followed by positive correlations while only positive correlations occur in correct responses (Cohen and van Gaal 2014). EEG signatures of errors also differ from those of conflict in correct responses because of their additional recruitment of delta power (Yordanova et al., 2004; Cohen and van Gaal, 2014) and phase-locked component (Trujillo and Allen, 2007; Munneke et al., 2015).

An intriguing point is that the phase of maximal power-RT correlation was not only subject-specific, but also condition-specific. This pattern, although robust in the results, is hard to interpret. Indeed, we don’t have any evidence that the different conflict conditions might re-align the phase of maximum correlation because stimuli in both studies were randomly displayed. Moreover, this was especially true for mixed correct trials, which were not even an experimental condition *per se* but a way to isolate the situations of high conflict based on the participants’ behavior (muscular activation of incorrect and correct responses). Further work will be needed to clarify this point.

### Midfrontal theta “clocks” large-scale cross-frequency coupling

A phase-dependency of a brain-behavior correlation suggests that certain computations are more likely to occur at certain theta phases. An ideal analysis to evaluate this hypothesis is to test for phase-specific activation of cell assemblies. In lieu of such an approach, however, we turned to phase-amplitude coupling, with the hypothesis that the behaviorally-optimal theta phase should also predict activity represented in higher frequencies. We applied a recently developed multivariate phase-amplitude coupling method that allowed us to differentiate between topographies of the phase-providing and the power-responding networks (Cohen 2017a). This analysis revealed that cross-frequency interactions were most pronounced in a fronto-parietal topography in the Simon task, and a fronto-lateral topography in the Eriksen flanker task. These topographical differences may be attributed to differences in the source of response conflict in these two tasks: although both tasks generate conflict as incongruency in response hands, *spatial* incongruity elicits conflict in the Simon task, whereas stimulus *identity* incongruency elicits conflict in the Eriksen flanker task. Such task differences have been shown to elicit slight differences in temporal and spatial dynamics of the brain activity (Frühholz et al., 2011; Nigbur et al., 2011).

Importantly, the pattern of theta-phase-locked time-frequency power looked remarkably similar despite the topographical idiosyncrasies, suggesting that the conflict-related theta rhythm provides a fundamental organizational principle that is used in similar ways by potentially distinct task-specific networks. In particular, the relevant theta phase was associated with power bursts in the beta band. This pattern was consistent across both datasets and was robust to two distinct statistical evaluation methods (permutation testing and cross-validation). Beta oscillatory activity has consistently been associated with motor control (Kilavik et al., 2013; Neuper et al., 2006), notably because of the exaggerated beta activity in Parkinson’s disease (Oswal et al., 2013). Motor-related beta is centered around the motor cortex and emerges as short-lived burst that are suggested to underlie response execution (Feingold et al., 2015). This suggests that the theta-phase-locked beta power bursts we found around motor areas relate to response execution. Moreover, we did not detect condition-specific modulations of multivariate phase-amplitude coupling. This also argues for a signature of a general response-related mechanism. On the other hand, condition differences might be too subtle or small-scale to be reliably detected with extracranial EEG. It is also possible that theta phase is regulating rhythmic fluctuations in attention through rhythmic enhancement of visual processing (Fiebelkorn et al., 2018), which might be important for task performance but equally prevalent across conditions.

### Guided source separation for guided network discovery

The kinds of findings reported here may be subtle (low signal-to-noise at the level of EEG), temporally transient, and spectrally narrowband. However, they are linearly distributed over many electrodes, which means that targeted multivariate analysis methods can help reveal patterns in the data that may be too mixed to uncover when analyzing only individual electrodes. We used a blend of hypothesis-driven (e.g., defining a component that maximizes theta activity) and data-driven (to define a component that maximizes specific theta phase activity) approaches in applying generalized eigendecomposition as spatial filters. This increased the signal-to-noise ratio and highlighted features of behaviorally-relevant brain activity that otherwise might have remained too subtle to be observed. We believe that these kinds of methods may be useful for elucidating subtle brain-behavior relationships in other contexts as well.

## Author contributions

**JD**: Conceptualization, Methodology, Formal analysis, Data curation, Writing - original draft, review & editing

**RG**: Conceptualization, Resources, Data curation, Writing - review & editing

**MXC**: Conceptualization, Resources, Methodology, Data curation, Writing - review & editing, Supervision

## Supplemental information

**Figure S1:**
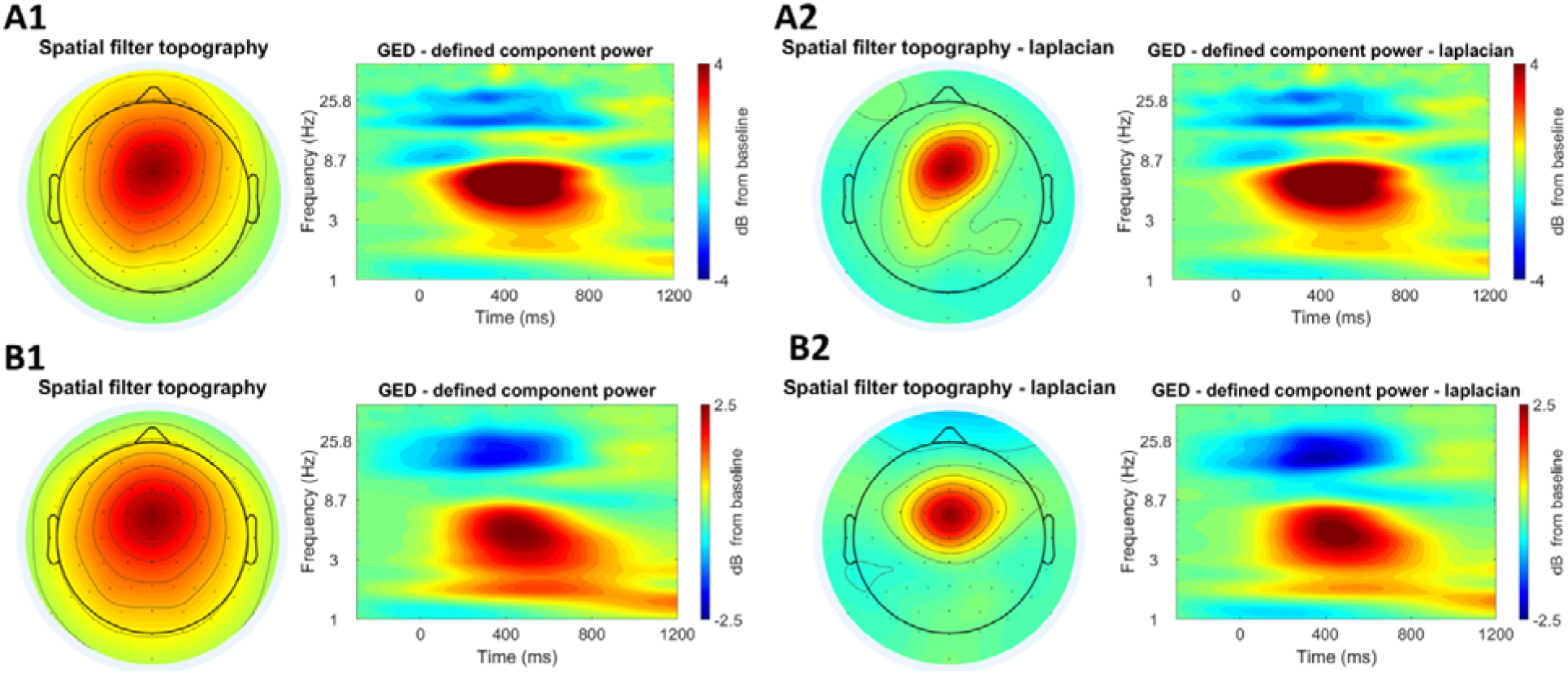
A1: Topographical map showing the activation pattern of the spatial filter designed by GED and the corresponding time-frequency power map for one subject in study 1 without the Laplacian spatial filter. A2: same plots with the Laplacian spatial filter. B1: Topographical map showing the activation pattern of the spatial filter designed by GED and the corresponding time-frequency power map averaged over all subjects in study 1 without the Laplacian spatial filter. B2: same plots with the Laplacian spatial filter.

**Figure S2:**
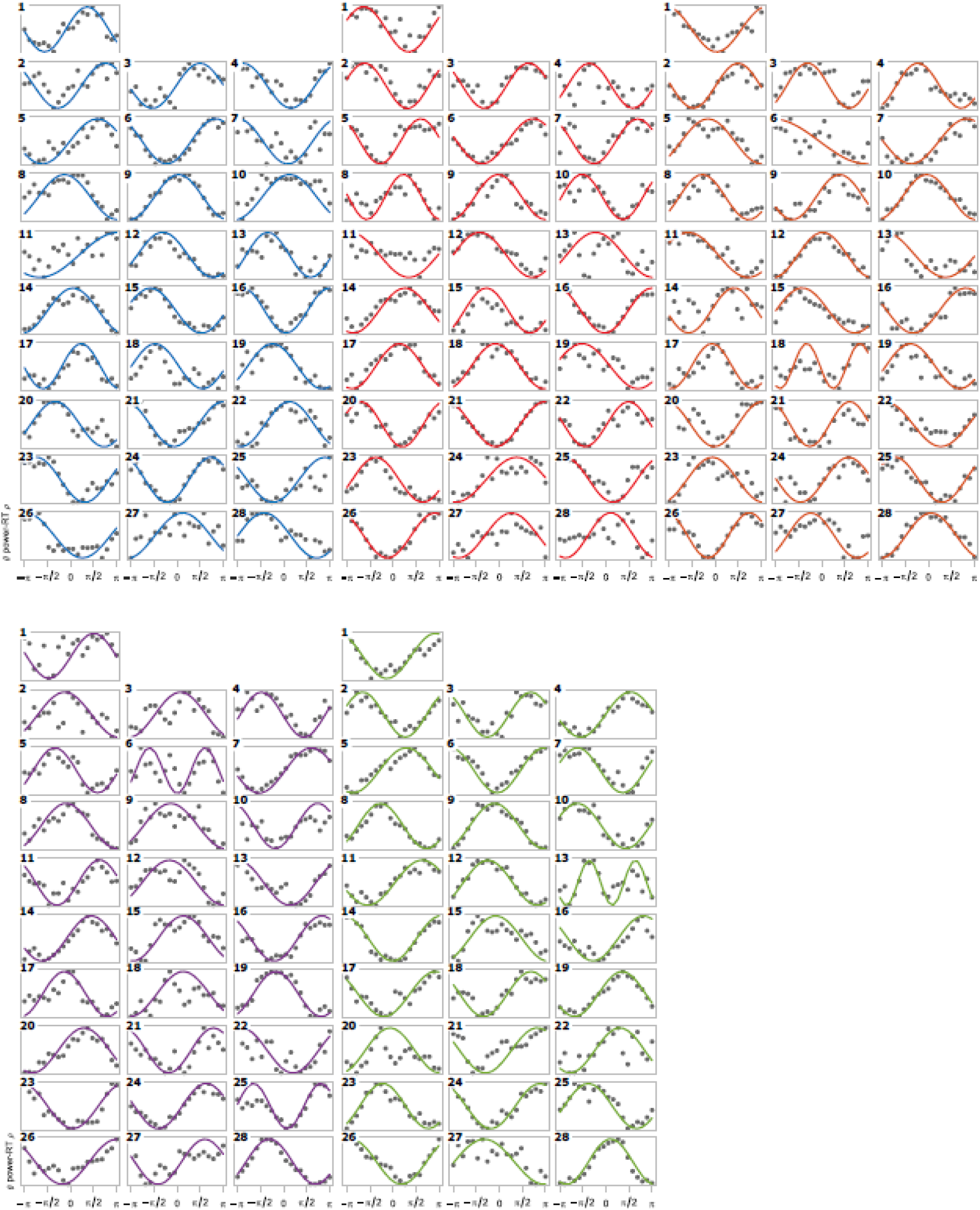
Sine fit plots of all subjects and in all conditions in study 1. The plots show theta power-RT correlation as a function of theta phase. Color coding of conditions is the same as in previous figures and goes from weaker to stronger conflict and errors from upper left to bottom right.

**Figure S3:**
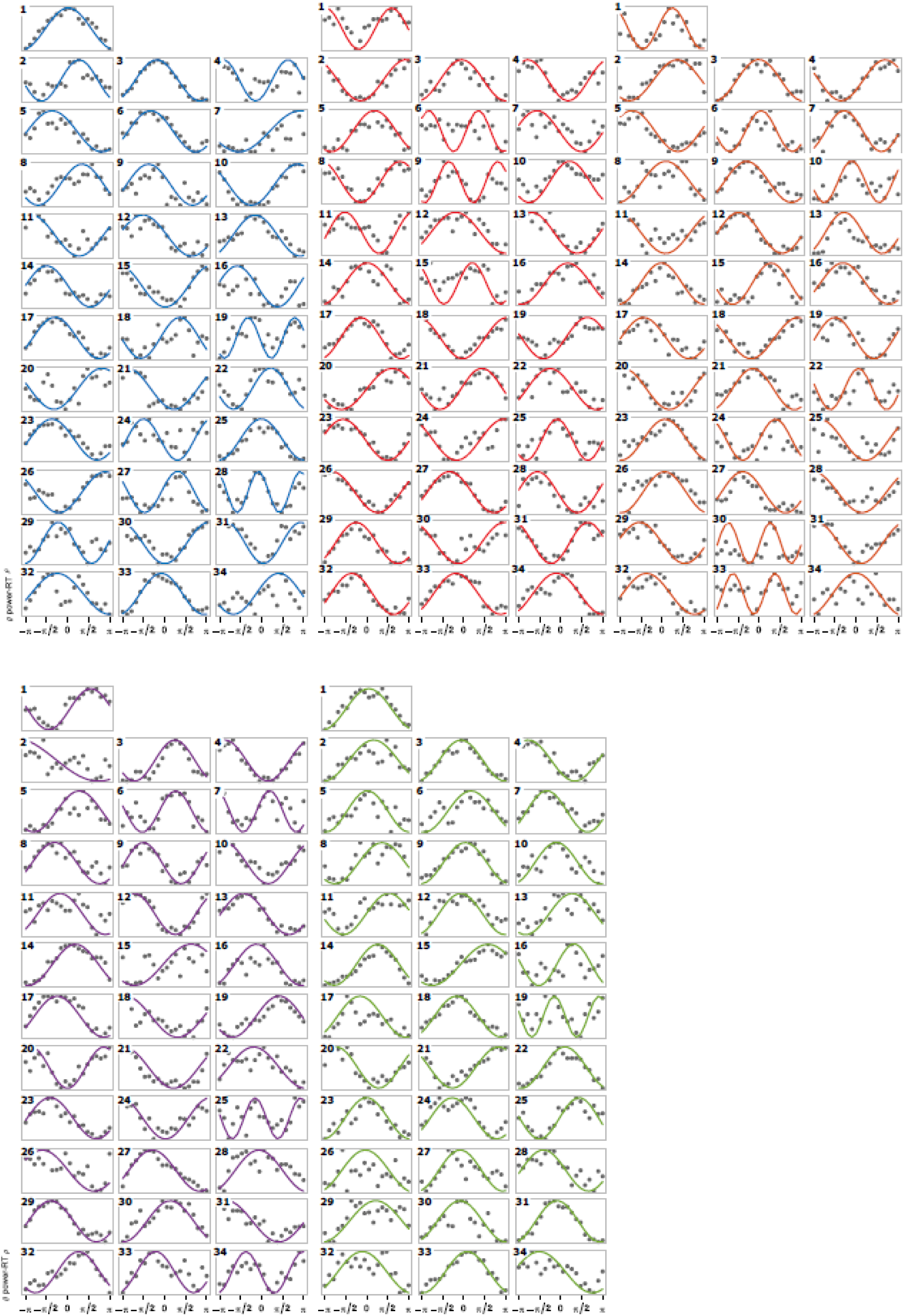
Sine fit plots of all subjects and in all conditions in study 2. The plots show theta power-RT correlation as a function of theta phase. Color coding of conditions is the same as in previous figures and goes from weaker to stronger conflict and errors from upper left to bottom right.

**Table S1:**
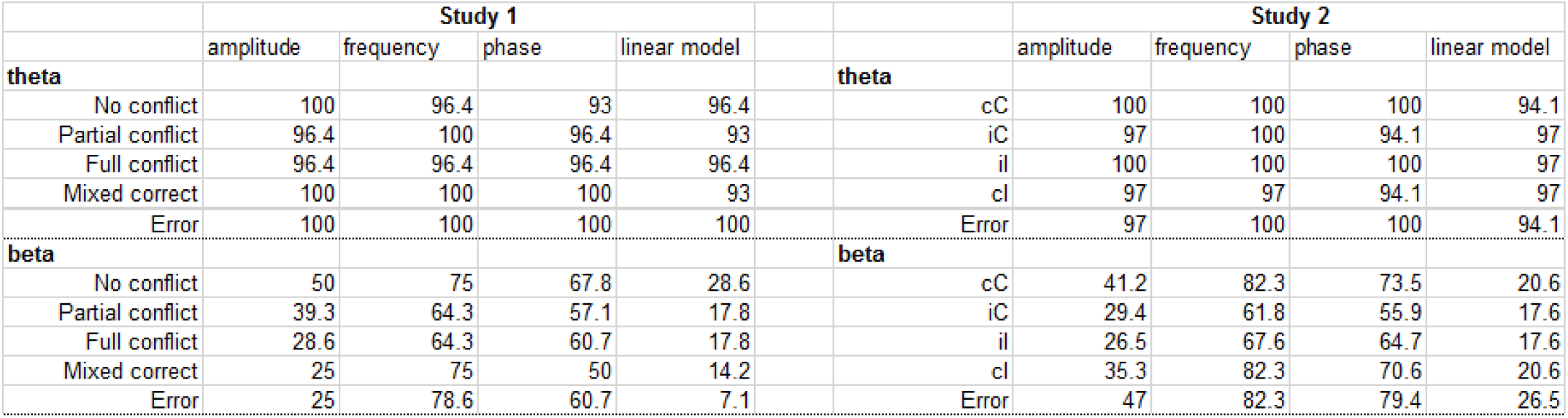
Overview of the sine fit results for the theta and beta analyses for study 1 and 2. Numbers show the percentage of significant parameters/model at p < 0.05 for each condition.

**Figure S4:**
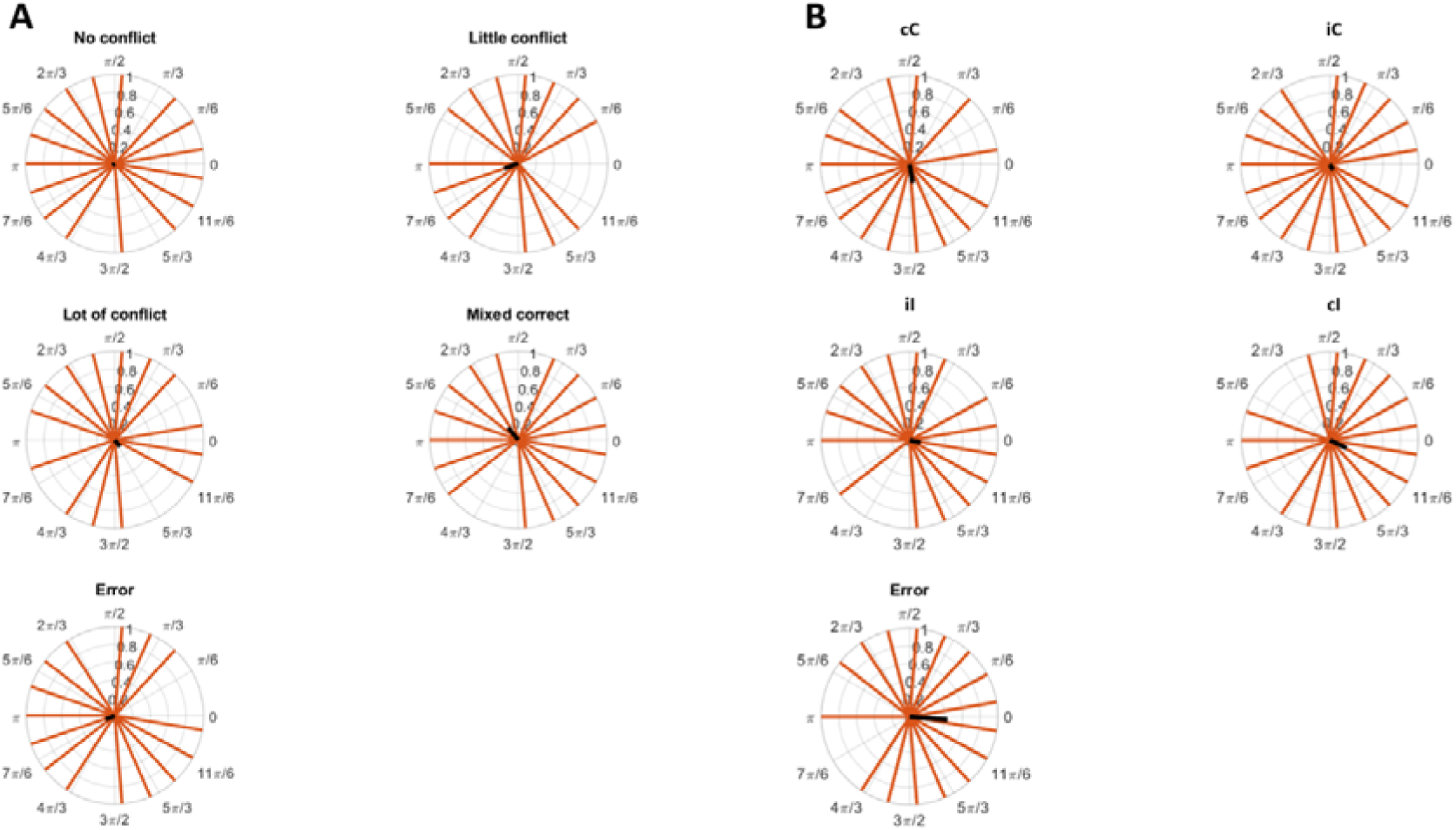
Clustering of phase of maximal power-RT correlation for each experimental condition for study 1 (A) and study 2 (B). Phase could only take 20 values since it was discretized in 20 bins. The thick black line represents the average complex vector, its angle represents the average phase angle, while its length indicates the size of clustering.

**Figure S5:**
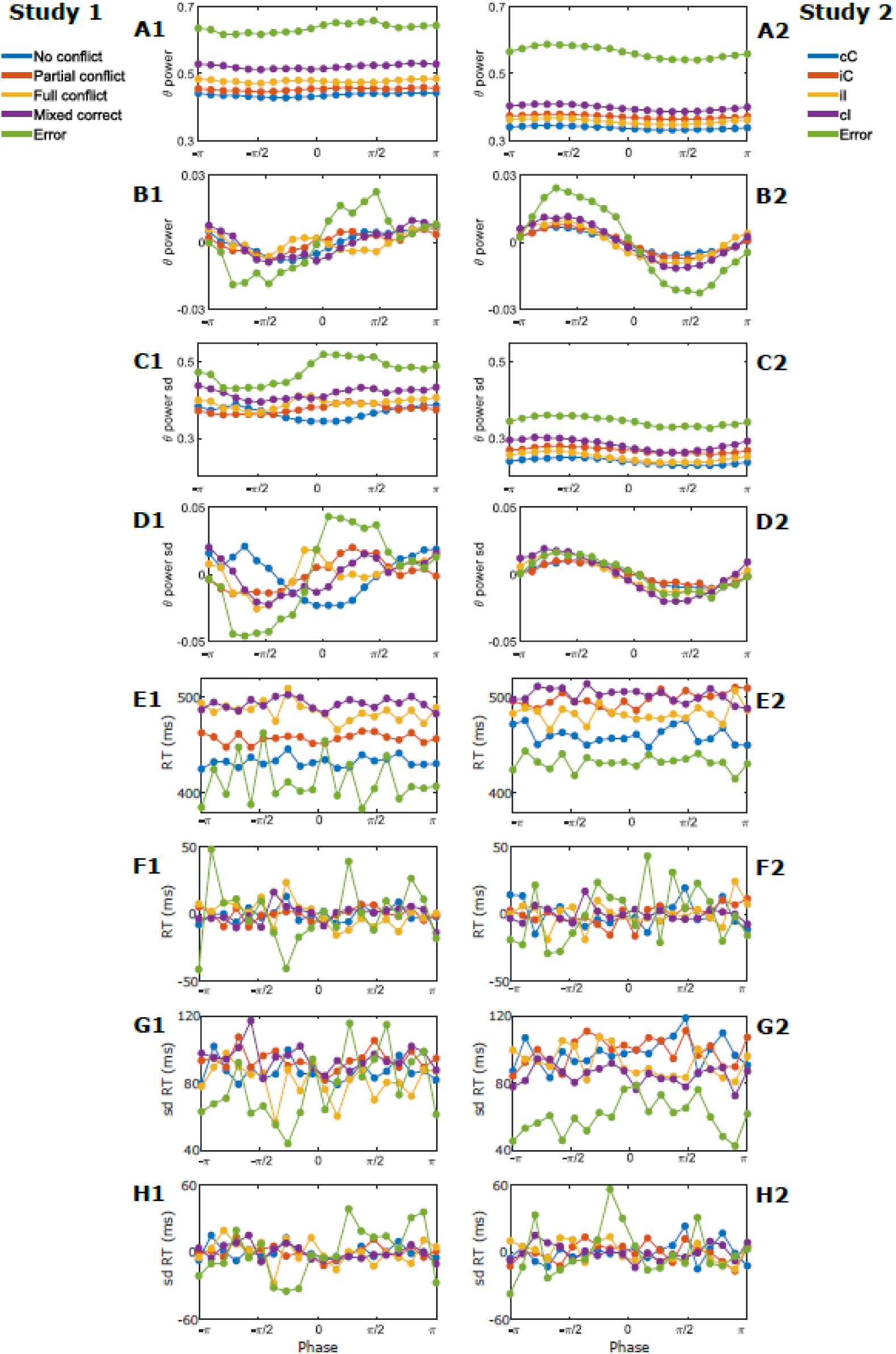
Theta phase resolved power and reaction time. Numbers 1 and 2 are for study 1 and 2 respectively. A and B show the phase resolved power and detrended power; C and D show the phase resolved standard deviation and the detrended standard deviation of power; E and F show the phase resolved RT and detrended RT; G and H show the phase resolved standard deviation and detrended standard deviation of RT.

**Figure S6:**
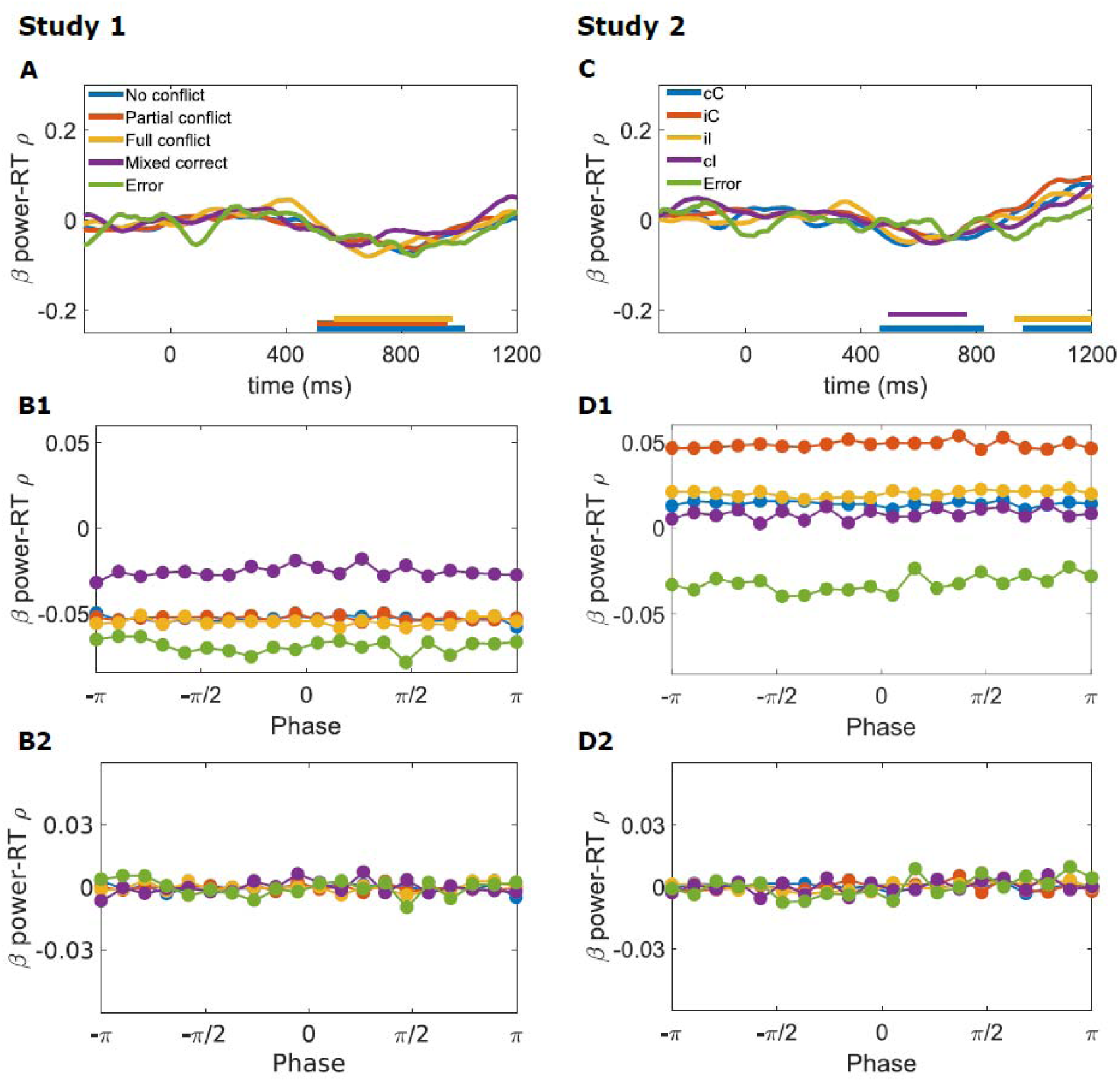
Time course of Spearman’s rho between beta power and RT averaged across subjects for each condition in study 1 (A) and 2 (C). Lower horizontal bars show time of rejection of symmetry around 0 at the group-level tested using permutation testing. B1 and B2: phase-resolved Spearman’s rho between beta power and RT. These plots show the average (B1) and detrended (B2) beta power-RT correlation coefficient according to beta phase. D1 and D2: same plots for study 2.

**Figure S7:**
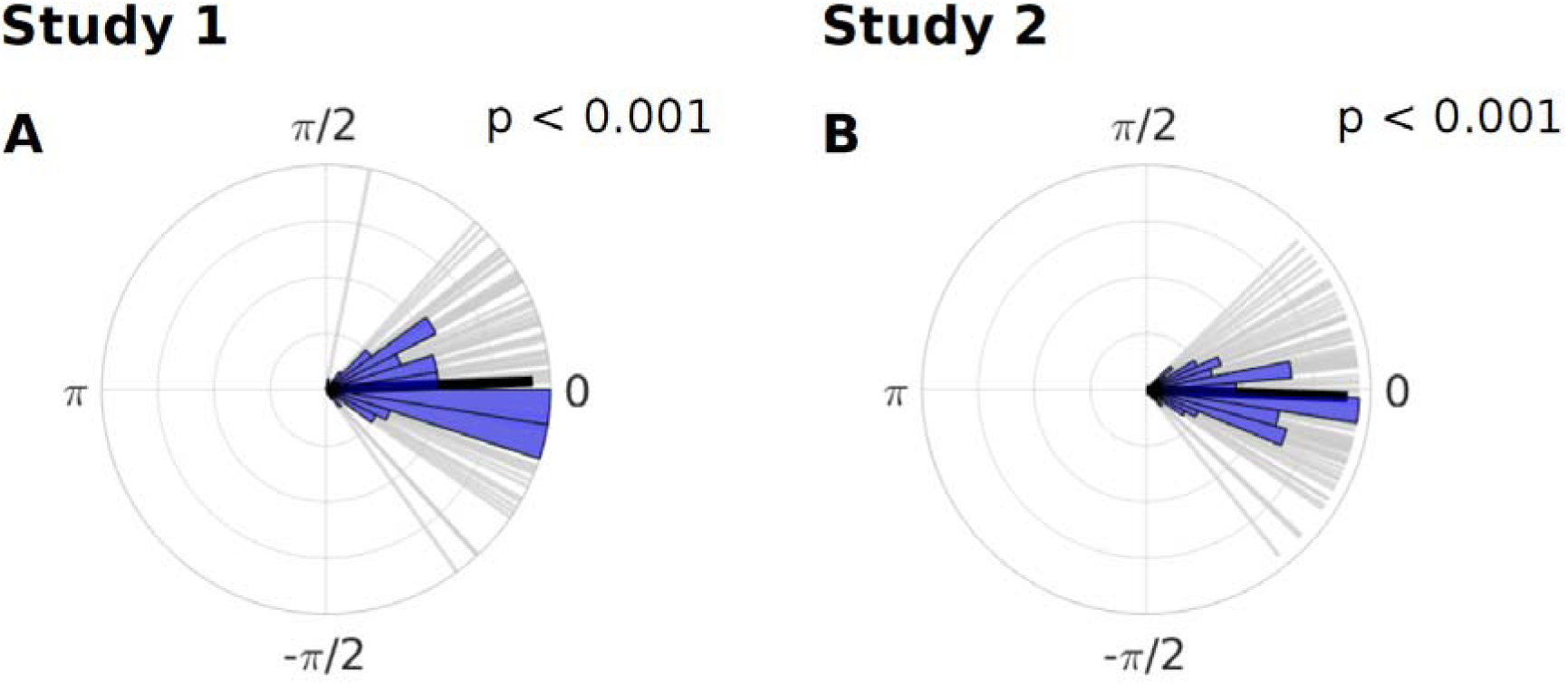
Clustering of phase of maximum correlation differences between two halves of data randomly selected over 100 iterations. Each grey bar represents the angle of the mean vector of phase preference differences between the two halves of the data over subjects for one iteration of random selection of data halves. Polar histograms in blue show the repartition of the phase differences mean vector angle. The black thick line represents the mean vector of all the 100 iterations of mean vector angles.

